# A simple method of suspending implant wear particles for more physiological modelling of cell-particle interactions *in vitro*

**DOI:** 10.1101/2021.01.05.425343

**Authors:** Christine Poon

## Abstract

Arthroplasty implants e.g. hip, knee, spinal disc sustain relatively high compressive loading and friction wear, which lead to the formation of wear particles or debris between articulating surfaces. Despite advances in orthopaedic materials and surface treatments, the production of wear debris from any part of a joint arthroplasty implant is currently unavoidable. Implant wear debris induces host immune responses and inflammation, which causes patient pain and ultimately implant failure through progressive inflammation-mediated osteolysis and implant loosening, where the severity and rate of periprosthetic osteolysis depends on the material and physicochemical characteristics of the wear particles. Evaluating the cytotoxicity of implant wear particles is important for regulatory approved clinical application of arthroplasty implants, as is the study of cell-particle response pathways. However, the wear particles of polymeric materials commonly used for arthroplasty implants tend to float when placed in culture media, which limits their contact with cell cultures. This study reports a simple means of suspending wear particles in liquid medium using sodium carboxymethyl cellulose (NaCMC) to provide a more realistic proxy of the interaction between cells and tissues to wear particles *in vivo*, which are free-floating in synovial fluid within the joint cavity. Low concentrations of NaCMC dissolved in culture medium were found to be effective for suspending polymeric wear particles. Such suspensions may be used as more physiologically-relevant means for testing cellular responses to implant wear debris, as well as studying the combinative effects of shear and wear particle abrasion on cells in a dynamic culture environments such as perfused tissue-on-chip devices.

## 1. Introduction

Arthroplasty implants are typically comprised of a metallic or ceramic piece and polymeric socket insert to respectively model the load bearing (bone) and shock absorbing (cartilage) components of the joint they are designed to replace. Polymers e.g. polyethylenes and polyetheretherketone (PEEK) are the most extensively used acetabular insert materials in total joint replacement prostheses due to their inertness and shock-absorbing properties. The other main classes of implant materials are metal alloys (e.g. titanium and cobalt chrome) and ceramics (e.g. zirconia and alumina) [1, 2]. Where the surfaces of two materials with different mechanical properties articulate, the softer material undergoes frictional wear and produces wear debris. As polymers are softer than metals and ceramics, wear particles tend to be generated from the polymeric component in any material combination involving polymers. Wear particles can also be produced from bone cement used to anchor the implant to healthy bone during surgical installation [3]. Arthroplasty implant wear debris causes periprosthetic inflammation, joint immobility, patient pain and mechanical instability of the joint [4]. It is understood that immune-mediated inflammatory responses to wear particles lead to osteolysis and aseptic loosening of the implant [5–8], which together reduces the life of the implant and necessitates revision surgery [6, 9, 10]. The size, geometry, load and intrinsic chemical reactivity of implant wear particles are known to determine the severity and rate of immune-mediated osteolysis [4, 11, 12]. In turn, these factors depend on the physical properties of the materials, wear mechanism (tribology) [13, 14], lubricant and patient condition [15]. Wear debris production can also be exacerbated by suboptimal positioning of the implant components during surgical implantation, as well as the size and geometry of implant components [16].

Characterizing wear debris is important for predicting wear rate, the development of more sophisticated wear-resistant materials, as well as for understanding the wear mechanisms of implant bearings in different loading and patient-specific scenarios. Furthermore, the role and material-specific physico-chemical properties of wear debris in provoking cellular and tissue responses that lead to osteolytic implant loosening require elucidation. *In vitro* studies of the effects of wear particles on cellular responses are valuable for preliminary evaluation of the cytotoxicity of periprosthetic implant materials. However, polymeric particles are light and tend to float, making it difficult to ensure sufficient contact of cell cultures with particles to elicit a biologically-relevant response. Several strategies to suspend wear particles within a culture environment have been described, including ultrasonic dispersion [17], emulsion in foetal bovine serum [18] and embedding particles in a low melting point agarose gel [19, 20]. However, these methods may either alter the size and physical properties of the particles by mechanical and thermal disruption or do not adequately model the cell-particle contact interactions that occur within the joint cavity and periprosthetic space, where wear particles free-float in synovial fluids prior to being phagocytosed by macrophages [21]. There is also a need for a synovial fluid analogue with suitable rheological properties for the study of implant wear mechanisms and the production of more clinically-relevant implant wear particles by wear simulators [22].

Here a simple method was devised to suspend polymeric wear particles in culture media for more effective biological testing by using sodium carboxymethyl cellulose (NaCMC) to thicken culture medium. Sodium carboxymethyl cellulose is an anionic, biologically inert, biodegradable, non-toxic, non-allergenic polymer that readily dissolves in water to form a hydrogel network. It is widely used as a thickening agent, emulsion stabiliser and viscosity modifier in the food and pharmaceutical industries as well as established applications as a hydrogel substrate in tissue engineering [23]. Furthermore, its polyelectrolyte structure and easily tuneable mechanical properties via concentration adjustment make NaCMC an ideal candidate material for generating sufficient viscosity in culture medium to suspend light, charged polymeric wear particles and heavier metallic particles. Importantly, like synovial fluid [24], CMC has thixotropic properties in aqueous media [25] which make it an ideal rheological analogue of synovial fluid for modelling wear particle suspension. In this work, NaCMC was found to easily dissolve in culture medium (RPMI-1640). Low concentrations (0.5-1.5%) of NaCMC were effective for achieving a homogeneous suspension of clinically-relevant volumes of polyetheretherketone (PEEK) particles by simple vortexing, negating the need for ultrasonic mixing or solvents.

## 2. Materials and Methods

### 2.1 Materials

NaCMC (MW 90,000) was purchased from Sigma Aldrich and RPMI-1640 culture medium was purchased from Thermo Fisher Scientific. Polyetheretherketone (PEEK) particles produced by a pin-on-disc wear simulator were supplied by [the University of Leeds, UK (source and material to be verified)].

### 2.2 Method

A 3% (w/v) stock NaCMC solution was made up by adding dry NaCMC to 50 mL of RPMI-1640 medium in a 50 mL capped tube. The tube of NaCMC-medium solution was placed on a shaker for 5 minutes, then kept refrigerated for 48 hours to allow the NaCMC to hydrate and dissolve. The tube was inverted and shaken for 3 minutes to ensure complete dissolution of the NaCMC, then further inverted and shaken several times prior to extracting volumes for serial dilution. Dilutions of 1.5, 1 and 0.5% NaCMC were made up with fresh RPMI-1640 medium in 10 mL capped tubes.

As the 3% NaCMC solution was found to be highly viscous, PEEK particles were added to the 1.5% dilution for preliminary proof of concept. A volume of approximately 25 mm^3^ of dry wear particles was then added to the tube of 1.5% w/v NaCMC RPMI-1640 solution and an equivalent volume of stock RPMI-1640 medium for comparison. The samples were vortexed for 3 minutes. The particle suspensions were then kept refrigerated and observed over 24, 48 and 72 hour increments, then diluted with fresh RPMI-1640 medium to 1 and 0.5% (w/v NaCMC) to determine whether particles remained suspended at lower concentrations of NaCMC. N.B. particles could also be added to the diluted NaCMC solutions. The method described here was carried out to determine the efficacy of serially diluting a concentrated particle suspension for dose-response studies.

## 3. Results and Discussion

NaCMC readily and easily dissolved in culture medium without the need for mechanical agitation by allowing for sufficient hydration time. Magnetic stirrers may be used should samples be required at shorter notice, otherwise stock NaCMC volumes can be made up in complete media and stored within a brief window of time in advance of biological studies. The initial concentration of NaCMC investigated (1.5% w/v) was found to enable effective incorporation and suspension of dry polymeric wear particles by simple vortexing. From Figure 1, it can be clearly seen that a significant proportion of PEEK particles floated and were unevenly dispersed in RPMI-1640 medium, while particles were homogenously suspended in the medium with addition of NaCMC. Particles remained suspended in the 1.5% (w/v) NaCMC solution over at least 3 days as well as when the suspension was diluted to 0.5% (w/v) CMC. Lower concentrations may be similarly effective for suspending polymeric wear particles and would be of interest to investigate further. Higher concentrations of CMC are likely required to suspend heavier particles produced from metallic implant materials. It is recommended that serial dilutions e.g. in 0.1% increments be performed to determine the minimum concentration of NaCMC required to keep study-specific wear particles submerged, dispersed and free-floating in culture medium.

**Figure 1.**
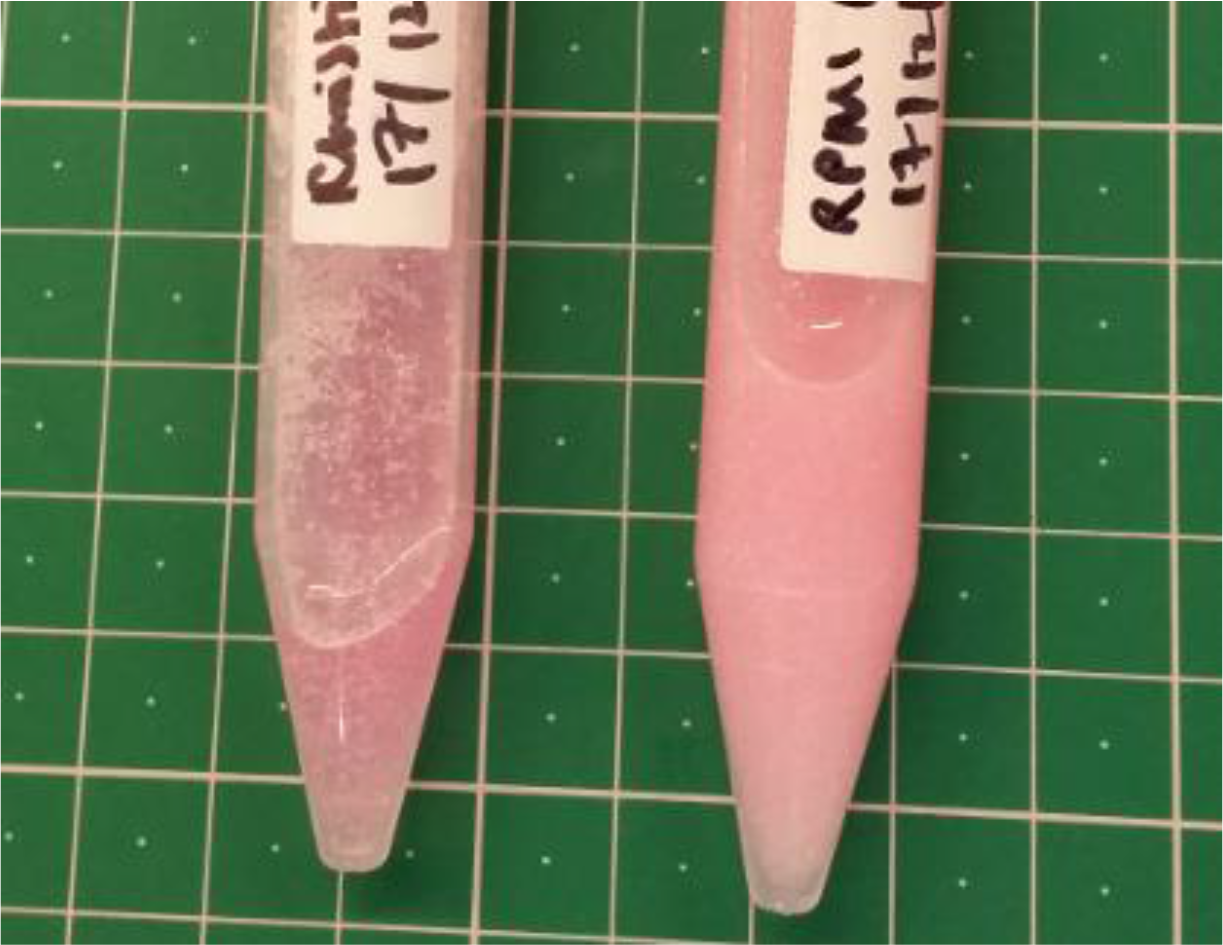
Left: PEEK particles (white) in RPMI-1640 medium with no added NaCMC. Right: PEEK particles were effectively suspended in 1.5% (w/v) NaCMC RMPI-1640 medium. Both samples imaged after 3 days and immediately after vortexing. N.B. The NaCMC-medium solution appears more opaque due to the presence of small air bubbles introduced during mixing.

A key advantage (and underlying rationale) behind the NaCMC-based suspension method reported here is that it provides a more physiological model of cell-particle interactions within the periprosthetic space, whether it be with macrophages, lymphocytes or surrounding tissues, compared to embedding particles in hydrogels such as agarose where there may be inconsistent and variable contact between cells and particles at the surface of the gel. These differences are illustrated in Figure 2. CMC solutions also have good optical clarity, which allows for imaging, and are compatible with a number of cell viability assay reagents. Another advantage of wear particle suspensions in liquid media thickened with NaCMC (ideally a rheological analogue of synovial fluid) is that they are theoretically compatible with mono or more complex 3D cocultures in gels e.g. culture of osteocytes or fibroblasts in a collagen gel similar to the protocol described in [26] (Figure 2B) and flow devices such as organ-on-chips, which would allow further study of the combinative effects of applied mechanical loading on particle distribution within 3D tissues and subsequent periprosthetic cellular responses. Systemic dissemination and the effects of wear particles on other tissues e.g. liver [27] can also be modelled and studied by applying a particle suspension to organ-on-chip devices. The technique is compatible with embedding particles in agarose gel, or for the study of particle infiltration into tissues where cells cultured in a gel layer then overlaid with the particle suspension (Figure 2B). Finally, while NaCMC was selected for this work, other celluloses and materials with established applications in tissue engineering such as alginate may also facilitate the suspension of wear particles in liquid culture media.

**Figure 2.**
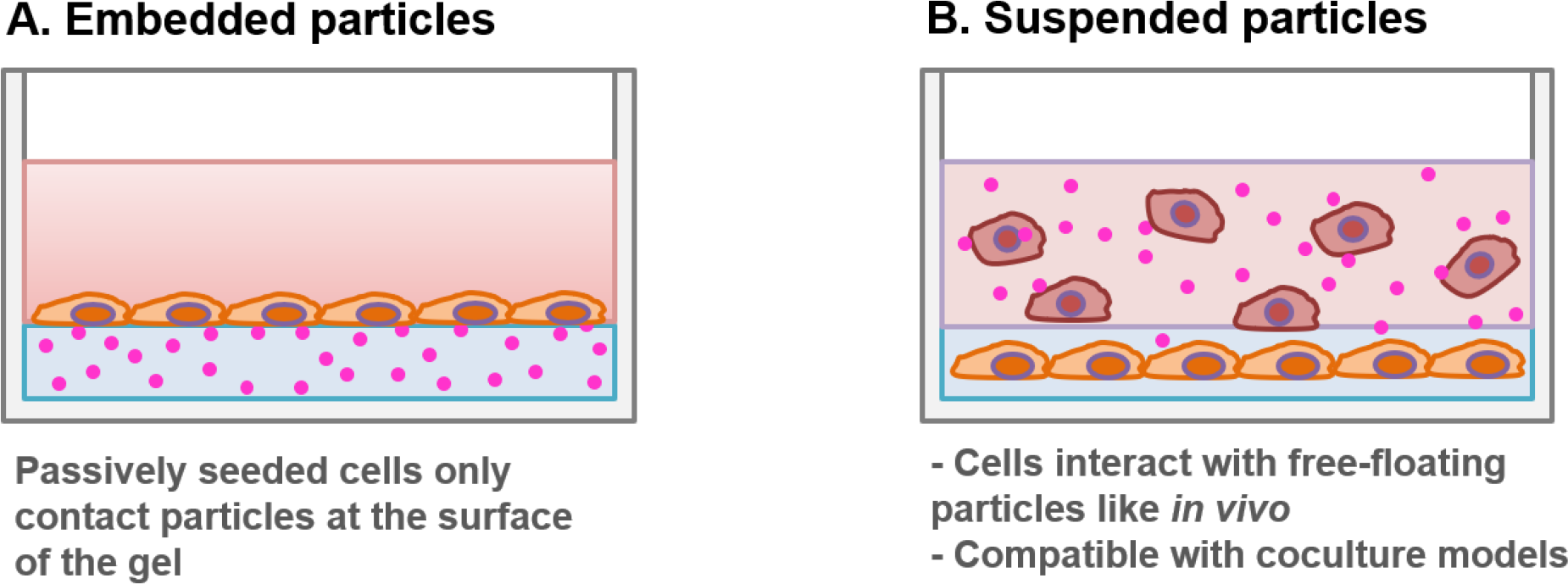
Comparison between embedding wear particles in agarose gel and suspension in media with low concentrations of added NaCMC

## Conclusions and Recommendations

This preliminary work demonstrated that low concentrations of NaCMC are effective for suspending light, polymeric wear particles in culture medium for studying and modelling cell-particle responses. The protocol can be readily adapted and modified for suspending particles of different densities by adjusting the concentration of NaCMC. It would be of interest to perform rheological characterisation to determine the concentration of NaCMC that would provide a physiological analogue for synovial fluid, keeping in mind that the molecular weight and degree of substitution of NaCMC, the quantity and type of added animal-derived sera for cell culture studies and batch and supplier variability would all affect rheological properties. While experimental parameters require optimisation as for the evaluation of any new model system, the effects of study-specific concentrations of NaCMC on cell viability can readily be investigated with colorimetric assays such as MTT. Lastly, a comparative study could be performed to investigate any differences between the inflammatory response of immune cells exposed to wear particles suspended by CMC in culture medium and those with the agarose overlay method.

## Acknowledgements

The author would like to thank the School of Biomedical Engineering, The University of Technology Sydney, Australia for supporting this work in terms of provision of materials and facilities.

## Author statement

The author confirms sole responsibility for the following: study conception and design, data collection, analysis and interpretation of results, manuscript preparation and ideation of further work.

